# The gastrointestinal development ‘parts list’: transcript profiling of embryonic gut development in wildtype and *Ret*-deficient mice

**DOI:** 10.1101/730010

**Authors:** Sumantra Chatterjee, Priyanka Nandakumar, Dallas R. Auer, Stacey B. Gabriel, Aravinda Chakravarti

**Affiliations:** Center for Complex Disease Genomics, McKusick-Nathans Institute of Genetic Medicine, Johns Hopkins University School of Medicine, Baltimore, MD 21205, USA; Center for Human Genetics and Genomics, New York University School of Medicine, New York, NY 10016; The Broad Institute, Cambridge, MA 02142, USA

**Author notes:** These authors contributed equally to the work (Sumantra Chatterjee, Priyanka Nandakumar). Send all correspondence to: Aravinda Chakravarti, Ph.D., Center for Human Genetics & Genomics New York University School of Medicine 435 East 30^th^ Street, Room 802/3, New York, NY 10016, E.

**Keywords:** Gut development, RNA-seq, *Ret*, non-autonomous cell effects

## Abstract

The development of the gut from endodermal tissue to an organ with multiple distinct structures and functions occurs over a prolonged time during embryonic days E10.5-E14.5 in the mouse. During this process, one major event is innervation of the gut by enteric neural crest cells (ENCC) to establish the enteric nervous system (ENS). To understand the molecular processes underpinning gut and ENS development, we generated RNA-seq profiles from wildtype mouse guts at E10.5, E12.5 and E14.5 from both sexes. We also generated these profiles from homozygous *Ret* null embryos, a model for Hirschsprung disease (HSCR), in whom the ENS is absent. These data reveal four major features: (1) between E10.5 to E14.5 the developmental genetic programs change from expression of major transcription factors (TF) and its modifiers to genes controlling tissue (epithelium, muscle, endothelium) specialization; (2) the major effect of *Ret* is not only on ENCC differentiation to enteric neurons but also on the enteric mesenchyme and epithelium; (3) a muscle genetic program exerts significant effects on ENS development, and (4) sex differences in gut development profiles are minor. The genetic programs identified, and their changes across development, suggests that both cell autonomous and non-autonomous factors, and interactions between the different developing gut tissues, are important for normal ENS development and its disorders.

**Significance statement:** The mammalian gut is a complex set of tissues formed during development by orchestrating the timing of expression of many genes. Here we uncover the identity of these genes, their pathways and how they change during gut organogenesis. We used RNA-seq profiling in the wildtype mouse gut in both sexes during development (E10.5 - E14.5), as well as in a *Ret* null mouse, a model of Hirschsprung disease (HSCR). These studies have allowed us to expand the universe of genes and developmental processes that contribute to enteric neuronal innervation and to its dysregulation in disease.

## Introduction

Vertebrate organogenesis involves highly regulated and evolutionarily conserved processes leading to the programmed differentiation of diverse cell types and their integration into a tissue or organ. The pathways involved are genetically programmed with genes expressed in precise spatial and temporal patterns and regulated by feedforward and feedback mechanisms, called gene regulatory networks (GRN) (1, 2). GRNs are modular, comprised of a small number of sub-circuit classes, conserved across species, and provide systems-level views of organogenesis (1, 2). They can also be the genetic basis for developmental disorders (3), as for Hirschsprung disease (HSCR, congenital aganglionosis), a multifactorial disorder of gastrointestinal development in which the ENS fails to develop (4).

The mammalian gastrointestinal tract develops from a tube to an organ comprising at least six well-characterized cell and tissue types (epithelial, smooth muscle, vascular, neuron, glia and extracellular matrix) that provide its major barrier function and its integrated physiology involving absorption, secretion and motility (5). The ontogeny of the gut involves cells that arise step-wise from multiple regimens of differentiation, being dependent on region-specific interactions between the endoderm-derived epithelium and the mesoderm-derived mesenchyme. Signaling between these two layers is critical for the differentiation and apoptosis of both epithelial and mesenchymal cells, and their subsequent homeostasis. The major developmental event in completion of early gut development is the migration of neural crest cells (NCC) into the gut, and their differentiation into enteric neuroblasts (ENCC) and, subsequently, enteric neurons and glia. These neuroblasts colonize the gut mesenchyme to form two neuronal networks, the myenteric (Auerbach’s) plexus, between the longitudinal and circular muscles, and, the submucosal (Messner’s) plexus, between the circular muscle and the submucosal layer. The myenteric and submucoal plexuses provide motor innervation to both muscular layers of the gut, and, secretomotor innervation of the mucosa nearest the lumen of the gut, respectively (6).

The many stages of gut development require numerous initiating signaling events activating TFs targeting diverse genes and pathways varying across development (7, 8). To improve our understanding of this process, we conducted gene expression profiling across gut development in wildtype mice. We also examined a mouse model of Hirschsprung disease (HSCR) which are null homozygotes for *Ret* (9, 10), the major gene for this disorder (4). *Ret* encodes a receptor tyrosine kinase (RTK) controlling ENCC differentiation and their migration through the gut mesenchyme. RET signaling is mediated by binding of a group of soluble proteins of the glial cell line-derived neurotrophic factor (GDNF) family ligands. RET does not directly bind to its ligand, but requires an additional co-receptor, one of four GDNF family receptor-α (GFRα) members (11). Mutations leading to Hirschsprung disease has been reported in many of these genes, highlighting the importance of this GRN in disease (12). *Ret* null mice exhibit complete aganglionosis with transcriptional changes in many of these genes (4), making it an ideal model to study how development is compromised in HSCR (10, 13).

Today, comprehensive transcriptome-wide studies allow the creation of a gene atlas from small amounts of developing embryonic tissue and identification of weakly expressing transcripts (14–16). Prior microarray studies of wild type and *Ret*-deficient mouse guts late in gut development (E15.5) has shown the dysregulation of many genes consequent to loss of *Ret* expression, including signaling molecules and TFs (17). However, we wanted to study the early effects of *Ret* loss uncompromised by secondary and downstream consequences. Here, we characterize the gut developmental ‘parts list’ as a function of developmental time (E10.5, E12.5, E14.5), sex (male, female) and *Ret* genotype (wildtype, homozygous null) to characterize its normal and HSCR genetic programs in development. We focus on the time period when the gut is differentiating into a mature organ and NCCs are undergoing migration, proliferation and maturation to ENCCs and enteric neurons (18). We identify bursts of activity of the activation of specific TFs prior to the differentiation and establishment of each major cell type and specialized tissue, and therefore its major regulators. We also demonstrate the dramatic effect of loss of *Ret* activity, on all major gut cell types. The effect of the gut mesenchyme on ENS development is well known; in distinction, we now show that loss of *Ret* identifies reverse crosstalk on the gut mesenchyme through multiple muscle-specific markers including Myogenin. Thus, both cell autonomous and non-autonomous genetic effects impact the ENS and its disorders. Finally, HSCR has a profound male sex bias, observed in appropriate mouse models (19), of unknown molecular etiology. This study shows that the gut transcriptional program is similar in males and females, with a few genes with specialized functions like cholesterol efflux and inflammatory response showing higher expression in females. Thus, additional mouse models need to be studied to understand the sex bias in HSCR.

## Results

We performed RNA-seq on dissected whole gut tissue at 3 embryonic stages during mouse development and assessed gene expression patterns using FPKM (Fragments Per Kilobase of transcript per Million mapped reads) values. Since there is no formal FPKM value that defines “expression,” we defined this quantity by qPCR analysis of selected genes. We randomly selected 7 genes with FPKM values between 0 and 6, at each developmental stage in the wild type male embryonic gut, on the same RNA samples used for RNA-seq. These results showed that in independent experiments using 40 qPCR cycles, genes with FPKM < 5 consistently had Ct values > 30 (**Figure 1**); thus, FPKM >5 was used as the expression threshold. We identified 9,436 and 9,432 genes expressed in the E10.5 wildtype male and female, respectively, or ~41% of the total transcriptome of 23,235 genes assayed. At subsequent times, only a small ~3% increase in the numbers of transcribed genes was noted, 10,143 (male) and 10,180 (females) genes at E12.5, and 10,216 (males) and 10,293 (females) genes at E14.5. Thus, increased developmental complexity is not associated with large increases in gene numbers *per se*.

**Figure 1:**
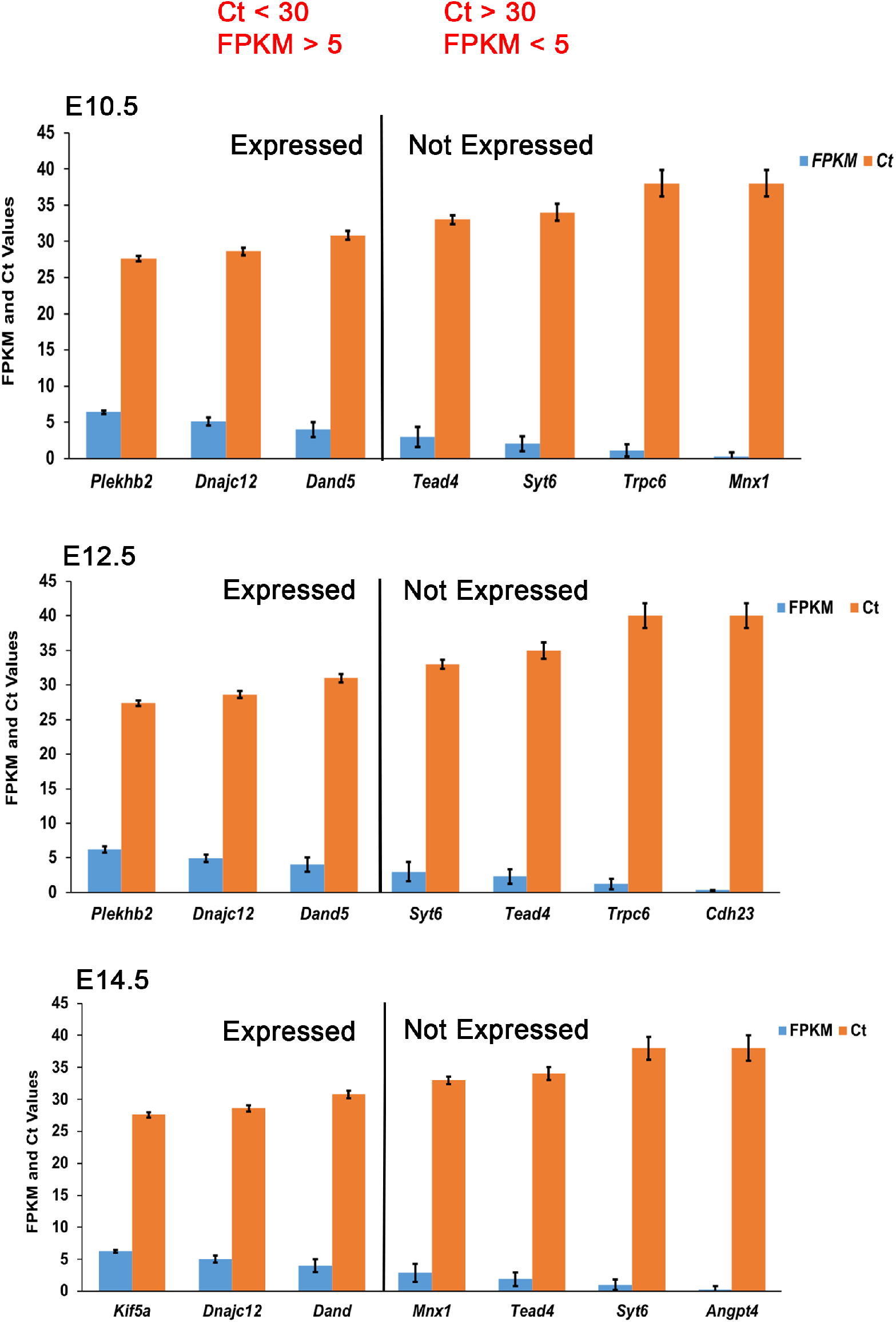
Gene expression in the embryonic mouse gut. Comparisons of FPKM from RNA-seq and threshold cycle (Ct) values after 40 cycles of qPCR of wildtype embryonic male mouse gut cDNA isolated at the developmental stages of E10.5, E12.5 and E14.5. We set FPKM = 5 as the value considered for a gene to be expressed; error bars are the standard errors of the mean for 3 biological replicates each.

### Relative effects of time, genotype and sex on gut development

In the first analysis, we attempted to identify groups of co-expressed genes depending on developmental stage, sex and *Ret* genotype and their interactions. We used signed network analysis to construct these modules, using Weighted Gene Co-Expression Network Analysis (WGCNA) (20) on 9,327 genes with a mean FPKM ≥ 5 across all 36 samples, i.e., 3-time points, 2 sexes, 2 genotypes (*Ret* wild type and null), 3 replicates. Expression values were transformed to log_2_(FPKM+1). Our analysis grouped 7,793 genes into 18 modules (labeled by colors) (**Additional Data Table 1**). Next, we tested associations between the first principal component (the eigengene) of each module’s expression profile with each of five binary variables using linear regression (genotype, sex and three pairwise developmental time comparisons), under a Bonferroni correction adjusting for 18 modules (p<0.00277) (**Figure 2**).

**Figure 2:**
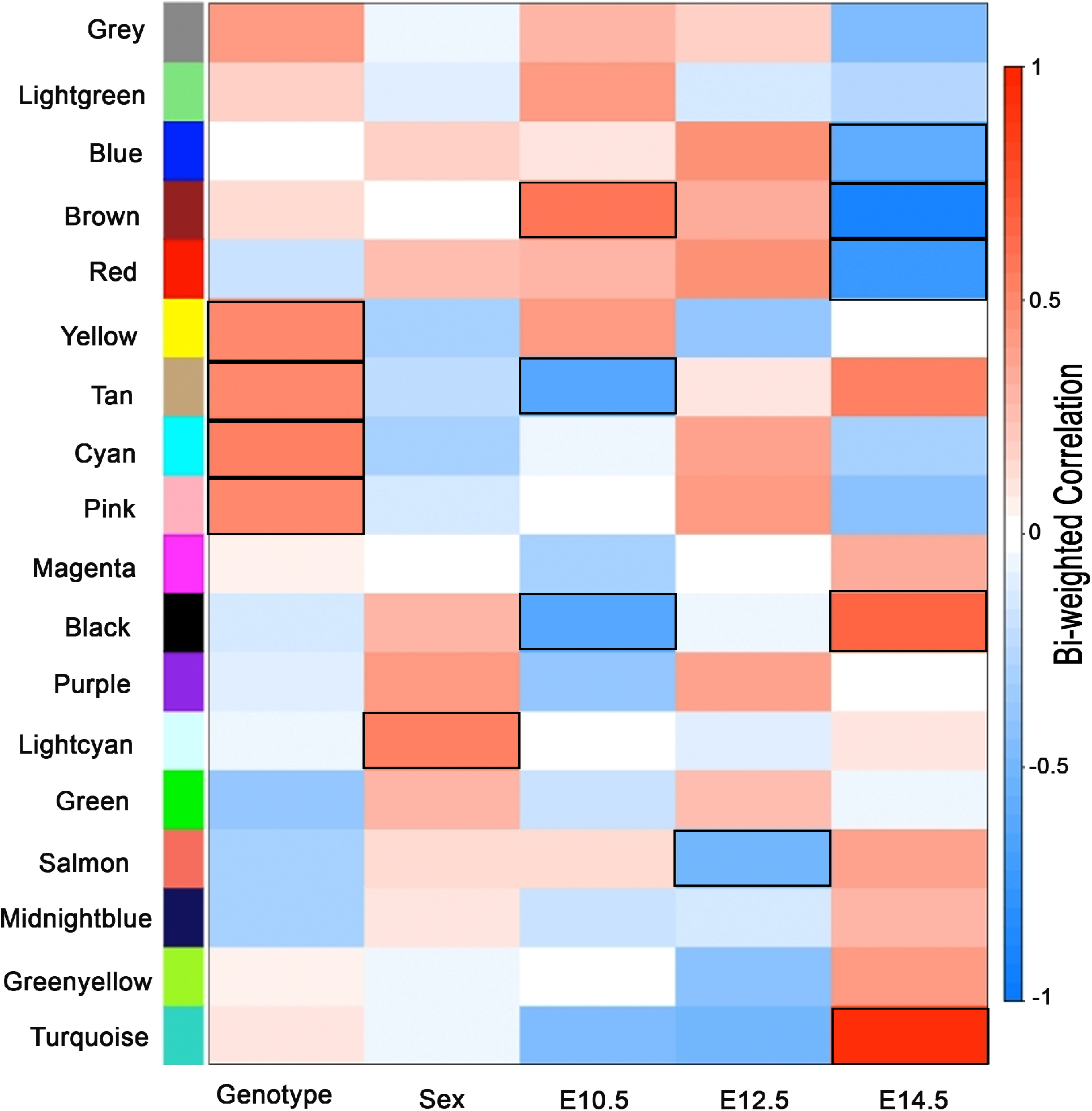
Weighted Gene Co-Expression Network Analysis (WGCNA). A correlation matrix using bi-weight mid-correlation in Weighted Gene Co-Expression Network Analysis (WGCNA) between 18 modules constructed from all expressed genes from 36 different conditions (3 time points, 2 sexes, 2 genotypes, 3 replicates). At a Bonferroni correction threshold adjusting for 18 modules (p<0.00277), the lightcyan module was found to be associated with sex (p=0.001), and, the tan (p= 0.0015), cyan (p= 0.0007), yellow (p=0.0026) and pink (p=0.0027) modules were found to be associated with genotype. The tan (p=6.18×10^−5)^, black (p=9.7×10^−5)^ and brown (p=0.0002) modules were associated with gene expression at E10.5 as compared to other time points. At E14.5, turquoise (p=5.2×10^−16)^, brown (p=4.6×10^−15)^, red (p=8.8×10^− 8)^, black (p=1.4×10^−5)^ and blue (p=2.7 x10^−4)^ modules were associated in comparison to other time points; salmon (p=0.0022, 62 genes) was associated with E12.5 only. All the associated module for each condition has been highlighted.

First, we identified a small lightcyan module containing 43 genes (0.55% of all genes in modules) associated with sex (p=0.001) and four modules comprising 1,092 genes (14% of all genes in modules) associated with genotype:yellow (p=0.0026, 694 genes), pink (p=0.0027, 199 genes), tan (p = 0.0015, 139 genes) and cyan (p= 0.0007, 60 genes). Three modules comprising 1,644 genes (21% of all genes in modules) were associated with early gene expression (E10.5 versus other time points): brown (p=0.0002, 1,210 genes), black (p=9.7×10^−5^, 295 genes) and tan (p=6.18×10^−5^, 139 genes). Only one module, salmon (p=0.0022, 62 genes, 0.80% of all genes in modules) was associated with E12.5 (versus others); the later (E14.5 versus others) gene expression program was characterized by five modules and 5,512 genes (71% of all genes in modules): turquoise (p=5.2×10^−16^, 2,208 genes), blue (p=2.7 x10^−4^, 1,443 genes), brown (p=4.6×10^−15^, 1,210 genes), red (p=8.8×10^−8^, 356 genes), and black (p=1.4×10^−5^, 295 genes). Note that by virtue of their reciprocal definition the black and brown modules are associated with both early and late development: the brown/black module contains genes expressed significantly higher/lower early/late in development.

In quantitative terms, thus, the greatest gene expression changes in gut development were across developmental time (21% and 71% of genes in early versus late programs), followed by *Ret* genotype (14%) together with a miniscule effect of sex (0.55%). In other words, the developmental parts list is skewed towards later use in development with differentiation into specialized structures and organ growth; early development is characterized by a differentiation program necessary to make the relevant cell types for later use. Thus, to understand the biological roles of these co-expressed genes in the various modules we performed gene ontology (GO) annotation using DAVID (https://david.ncifcrf.gov/) (21) and assessed the statistical significance of the grouped annotated functions at a false discovery rate (FDR) of 1%.

### Genes and pathways associated with developmental time, Ret and sex

Three modules were associated with developmental time. The brown module, with significantly higher expression in E10.5 versus other times, is enriched for genes controlling transcriptional modulation (*Med24, Med22, Nono etc.*), RNA processing (*Crnkl1, Supt6, U2af2 etc.*) and transcription factors (*Foxo4, Gli3, Sox11,* etc.) (**Figure 3**). In contrast, the black module comprises genes expressed at significantly lower levels in E10.5 versus other times and whose activities control macromolecule and protein transport (see below). The tan module also contains genes with weak expression at E10.5 versus other times: although no functional annotations were significantly enriched, it includes genes of the canonical Wnt signaling pathway (*Sfrp5, Fzd5, Tax1bp3* etc.), transcriptional repressors (*Snai2, Limd1* etc.) and negative regulators of cell proliferation (*Ptprj, Raf1, Asph* etc.). Thus, transcription and cell proliferation are the primary processes underlying gut organogenesis at E10.5. At E12,5, only the salmon module is associated with gene expression and it contains genes for protein translation (*Dnajb11, Pdia6, Pdia4 etc.*), protein transport (*Tmed4, Tomm6, Mcfd2 etc.*) and genes involved in steroid metabolism and homeostasis (*Npc2, Lamtor1, Ehd1 etc.*). Interestingly these processes are still negatively correlated at this stage of gut development, similar to E10.5 (black module), highlighting the continuity of function or lack thereof of specific genes during gut development (**Figure 3**).

**Figure 3:**
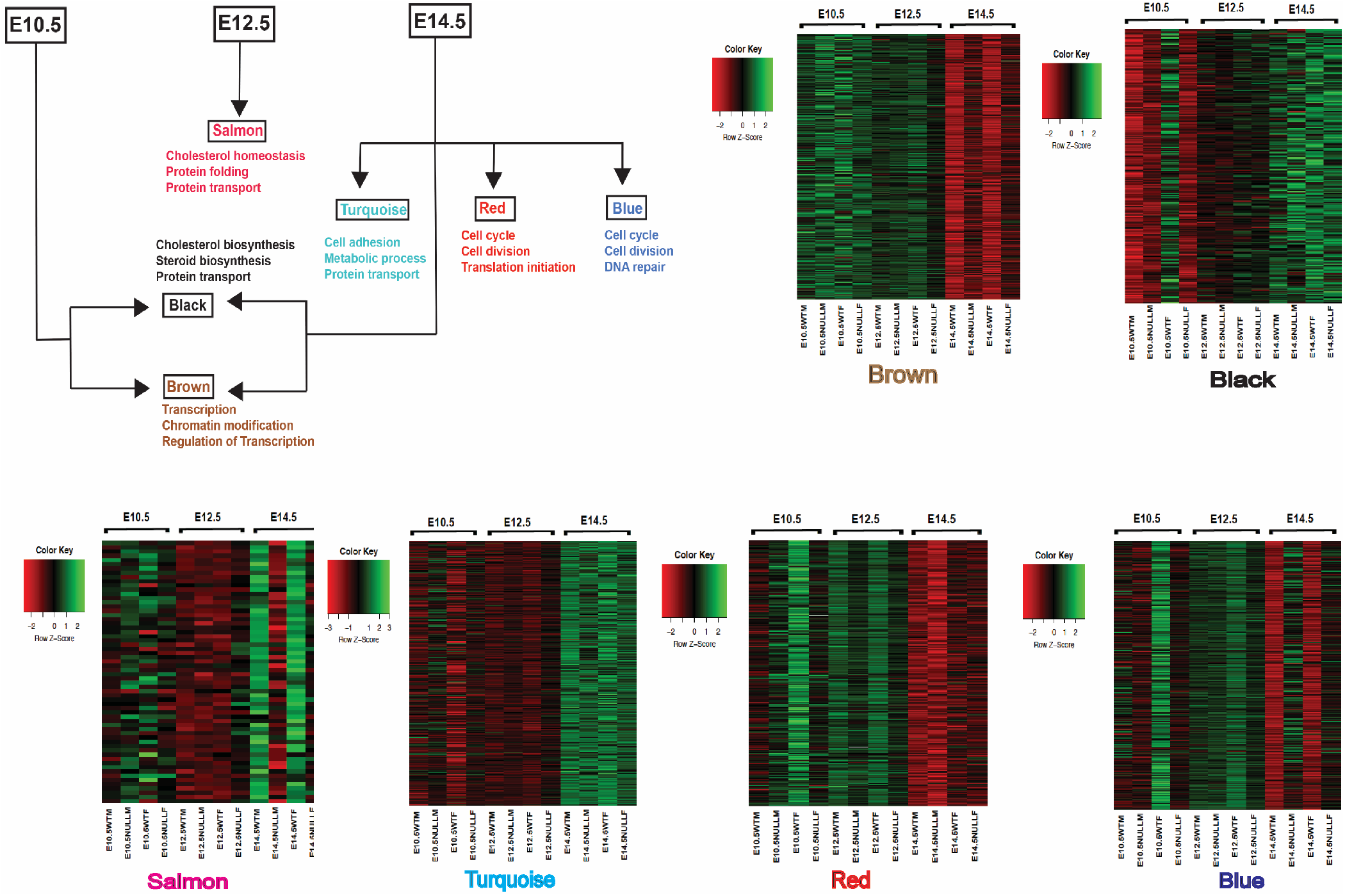
Biological processes affected by genes changing over developmental time. Various modules are associated with different developmental time points and shows a continuity of biological processes over time. The brown module is associated with E10.5 (early gut expressed genes) and the GO biological processes these genes are involved in include mostly transcription factors and chromatin modifiers. The Salmon module associated with E12.5 contain genes involved in protein folding and transport which are also seen in the Black and Turquoise modules, which are associated with E14.5. The Red and Blue module associated with E14.5 contain genes involved in the cell cycle, cell division, DNA repair and are weakly expressed at this stage when compared to early developmental stages. The corresponding heatmaps for all modules are also shown.

Two modules (black and turquoise) were associated with higher and three (red, blue and brown) with lower gene expression at E14.5, as compared to earlier stages. Genes in the black module are involved in transport of macromolecules, small molecules, ions/cellular components (*Uqcrfs1, Slc35a3, Gpr89, Hook1 etc.*) and steroid/cholesterol biosynthesis (*Cyp51, ACbd3, Sc5d etc.*), while turquoise module genes include those regulating metabolic processes in cell growth (*Ide, Ogdh, Fdft1 etc.*), as well as transport of macromolecules, small molecules, ions/cellular components *(Slc9a8, Atp1b1, Sec31a etc.*) and cell adhesion (*Lima1, Ldha, Tln1, etc.*). Thus, these later functions are involved with producing and transporting basic macromolecules necessary for survival, maturation and growth of the organ (**Figure 3**). Genes suppressed during this stage include genes for cell cycle progression (*Cks1b, Cetn3, Bex2 etc*), cell division (*Spc24, Ccne1, Kif2c etc.)*, RNA processing (*Raly, Ncbp2, Lsm6 etc.*) and translational initiation (*Eif3d, Eif4e, Eif3g etc.*) from the red module, and, genes controlling cell cycle progression (*Dbf4, Kntc1, Cdc16 etc.*), cell division (*Ccnt2, Cdk19, Tsg101 etc.*) and DNA repair (*Xrcc5, Xrcc2, Setx etc.*) from the blue module **(Figure 3**). Consequently, at E14.5, cell division is not as rapid as earlier in development and the tissue is reaching a morphological developmental equilibrium.

Beyond these qualitative changes, we also identified temporal shifts in the transcription of genes controlling specific gut processes such as cholesterol homeostasis and protein transport. These features are negatively correlated at E10.5 (black) and E12.5 (salmon) but positively correlated at E14.5 (black, turquoise). Understanding the roles of this temporal dynamic behavior in gut disorders may be important.

Gut motility is controlled by the ENS which monitors the state of the lumen and the gut wall by activating intrinsic reflexes that generate peristaltic movements, change blood flow, water and electrolyte secretions. The ENS arises from vagal NCCs that migrate into the foregut and then migrate caudally (18, 22) and then undergo fate transition first to ENCCs and then to enteric neurons. The receptor tyrosine kinase gene *Ret* is the earliest, major and key factor controlling ENCC differentiation, proliferation and migration. Therefore, it is no surprise that *RET* is the major gene for HSCR in humans with both rare coding and polymorphic enhancer variants (23–25). Further, *Ret* gene deletions in the mouse recapitulate most of the HSCR phenotype (6, 10, 13).

We identified gene modules that are *Ret*-genotype, wildtype and null homozygotes, dependent. Genes associated with the cyan module, with higher gene expression in *Ret* null versus wildtype homozygotes, are involved in neural crest cell differentiation *(Edn3, Isl1, Gdnf etc.*), enteric neuron differentiation (*Hdac5, Gfra1, Isl1 etc.*) and transcription factors and activators like *Tcf21, Ntf3, Foxf1 etc. (***Figure 4A**). These results are consistent with our earlier observations of *Gdnf* and *Gfra1* (4). In contrast, genes in the yellow module, with higher gene expression in *Ret* wildtype versus null homozygotes, comprise those involved in transcription and its modulation (*Zfp335, Med25, Zfp580 etc.*) and chromatin modifiers (*Ing4, Brd2, L3mbtl2 etc.*) (**Figure 4B**). Thus, *Ret* appears to have a significant effect on many early TFs and may explain why *Ret* mutations have such severe consequences. The pink module contains genes weakly expressed in wildtype, as compared to the Ret null homozygote, and includes genes for cell differentiation (*Wnt5a, Srpk2, Rbm45 etc.*) and nervous system development (*Sema5a, Itga8, Efna5 etc.*). These results show marginal statistical significance but suggests that certain differentiation and ENS genes may also be suppressed by *Ret* highlighting its balancing role in ENS development. Similarly, the genes in the tan module are also weakly expressed in wildtype and grouped primarily into broad categories like digestive tract development (*Fgfr2, Kit, Lgr4* etc.), negative regulation of cell proliferation (*Sfrp5, Ptprj, Raf1* etc.) and cytoslelekton reorganization (*Pak1, Flna, Parva* etc.). But like the pink module all functional groups show nominal significance.

**Figure 4:**
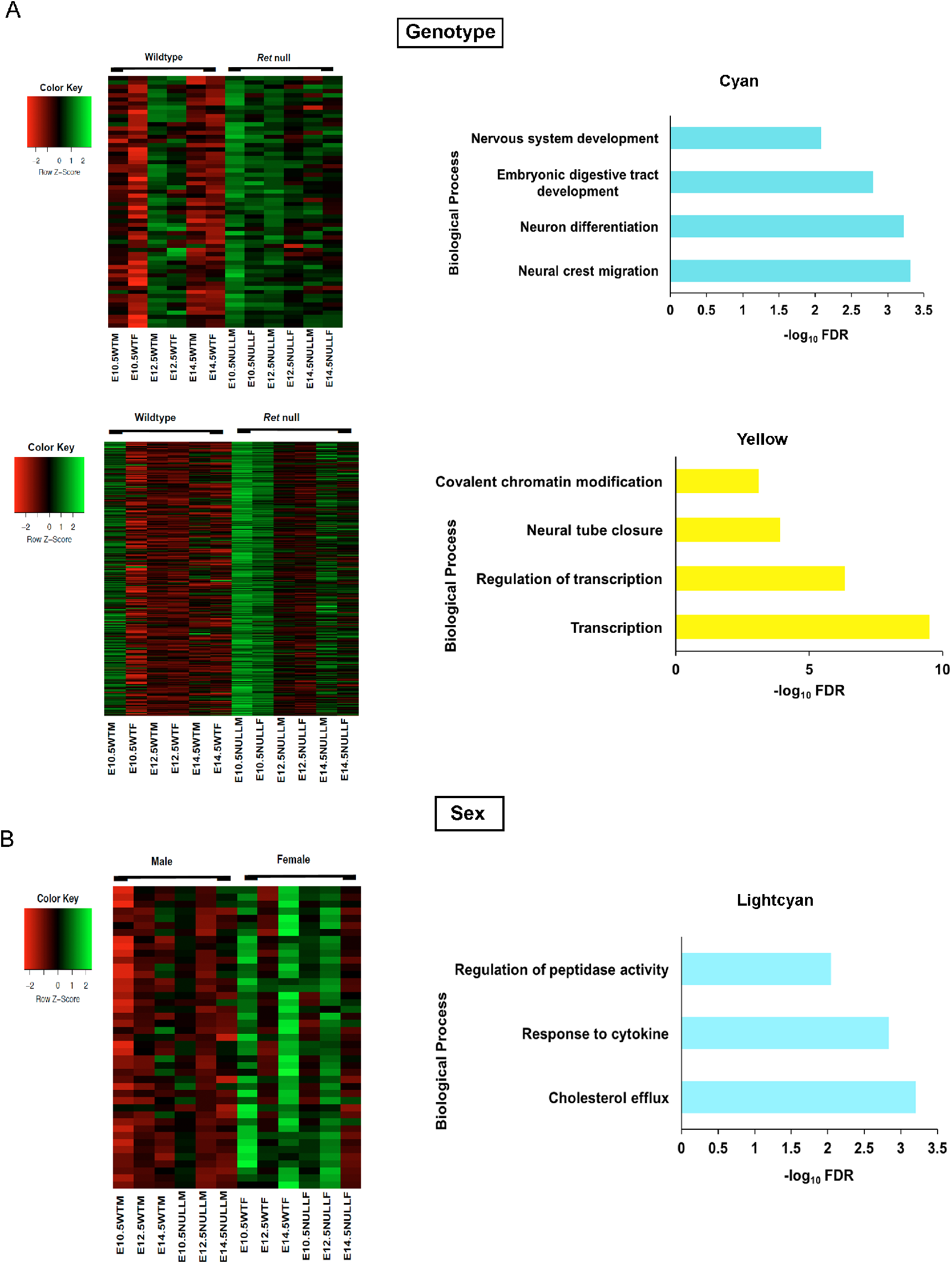
Biological processes affected by genes changing due to genotype and sex. (A) Heat map of GO biological processes for the two modules associated with genotype shows genes involved in neural crest migration and neuron division (Cyan), while (B) transcription and chromatin modifiers (yellow) are affected with loss of Ret transcription. (C) The lightcyan module consist of 43 genes for peptidase activity and cholesterol transport and is associated with sex.

We identified only one module (lightcyan) of genes significantly associated with sex, irrespective of genotype and developmental time. This set contains 43 genes broadly classified into those regulating cholesterol efflux (*Apoa2, Apoa1, Apoc1, etc.*) and regulation of peptidase activity (*Ambp, Serpina1b, Serpinf2 etc.*). Surprisingly, all showing higher expression in females than males (**Figure 4C**). Of these only *Cited1* is sex-linked showing that sex-linked dosage effects are not the major reason for these differences, suggesting hormonal or sex-biased epigenetic effects instead.

The miniscule sex-difference we identified, despite its prominence in developmental disorders of the gut, such as in HSCR, prompted us to look for such differences more carefully within each developmental stage. At E10.5, 270 genes show significant expression differences between wildtype males and females (**Figure S1A**), of which 157 and 113 show higher expression in males and females (**Additional Data Table 2**), respectively, a skew decidedly increased towards males (P = 0.007). Annotation analysis using DAVID showed that male enriched factors encoded genes for cell morphogenesis and cell motility and neurogenesis (**Figure S1B**). In contrast, the 113 female enriched factors encoded genes for hemostasis, vasculature development and blood coagulation (**Figure S1C**). Later at E12.5, 81 and 115 genes show higher expression in males and females, respectively, but, now, a skew decidedly favoring females (**Figure S1D and Additional Data Table 3**), a skew also observed at E14.5 where 133 and 169 show higher expression in males and females, respectively (**Figure S1G and Additional Data Table 4**). Interestingly, genes expressed significantly at a higher level in males are involved in specific morphological functions like neuronal, musculature and endothelial development (**Figure S1B, S1E and S1H)** while female enriched genes control specific physiological processes like response to wound healing, inflammatory response and response to hormonal stimulus (**Figure S1C, S1F and S1I).** Thus, sex differences do occur and occur differentially by functional categories. These results prompted us to search for genetic programs in gut development by more focused pairwise analyses of time and *Ret* genotype.

### Gene expression changes during normal gut development over developmental time

To understand stage-specific gut development we next compared gene expression changes between early E10.5 (early) versus E14.5 (late) stages in the wild type gut to discover 2,697 differentially expressed genes **(Figure 5A and Additional Data Table 5),** of which 1,227 showed significantly higher expression in E10.5. We performed GO annotation on only the subset of 316 (25.8%) of these genes with 2-fold or greater expression and identified 89 transcription factors (*Bach1, Gata2, Barx1* etc.), 73 genes for transcriptional control including various transcription factors like *Gli3, Hoxc6, Hoxc9* etc. and 29 for morphogenesis (*Crabp2, Prrx1, Col2a1*etc.) (**Figure 5B**). Conversely, of 1,470 genes showing significantly higher expression at E14.5, 1,003 genes (68.2%) with 2-fold and greater expression were enriched for biological processes like cell adhesion (76 genes which include *Cldn8, Cldn7, Aebp1* etc.), extracellular matrix organization (20 genes including *Olfml2b, Eln, Col3a* etc.), regulation of muscle contraction (44 genes including *Myl6, Tbx20, Myh7* etc.) and vasculature development (61 genes including *Cspg4, Anpep, Cav1* etc.) (**Figure 5C**). These results validate our WGCNA analysis that early gut morphogenesis is dictated by transcription factors whereas later development focuses on specifying specialized tissues. This analysis also identified the specific transcription factors which are candidate genes for genetic disorders of early gut development (see below).

**Figure 5:**
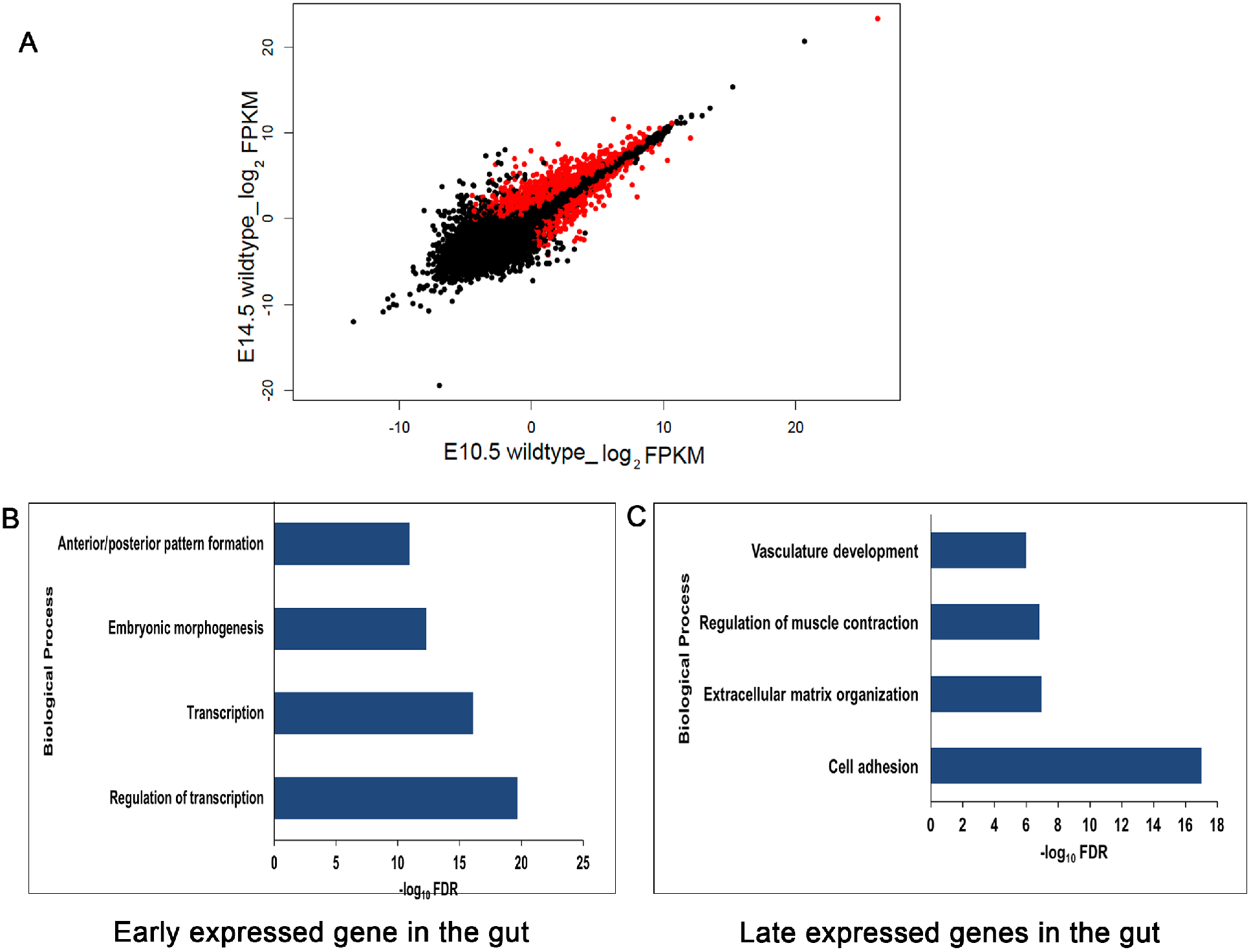
Temporal gene expression patterns in the embryonic mouse gut. (A) Scatter plot of log_2_ FPKM values of genes expressed in wildtype male embryos during early (E10.5) versus late (E14.5) gut development. (B) GO annotations of the transcriptome shows that transcription factors are highly expressed at E10.5, while genes controlling more specialized structures (e.g., vasculature) or processes (e.g., cell adhesion and cell signaling) are prominent at E14.5. Genes marked in red have statistically significant (q-value <0.01) expression differences between the two states indicated.

### Ret-dependent gene expression in the gut

Loss of *Ret* gene expression in the mouse gut has transcriptional consequences on many genes in the *Ret* GRN (4), as observed in our WGCNA analysis, but we sought to identify additional temporal patterns of gene expression changes triggered by *Ret* loss-of-function by detailed pairwise analysis. At E10.5, when *Ret* is normally expressed and is the earliest stage where reduction in the numbers of ENCCs and their failure to migrate can be demonstrated in *Ret* null homozygote mice (26), we identified 763 genes with significant expression differences (**Figure 6A and Additional Data Table 6)**, of which 521 were down-regulated (thus, *Ret*-activated) and 242 were up-regulated (thus, *Ret*-suppressed) in male *Ret* null embryonic guts as compared to wildtype. The *Ret*-activated genes broadly comprise three biological processes: neuronal development (Ret’s primary function), transcriptional regulation and pattern specification, primarily TFs (**Figure 6B)**. In contrast, the *Ret*-suppressed genes are involved in extracellular matrix organization and metabolic processes (**Figure 6C)**. At E12.5, there are 353 downregulated and 90 upregulated genes (**Figure 6D and Additional Data Table 7**), and, at E14.5, similarly, there are 325 down regulated and 111 up regulated genes (**Figure 6G and Additional Data Table 8**) in *Ret* null versus wildtype male embryos. The *Ret*-activated genes at E12.5 are involved in neurogenesis but, surprisingly, also genes for muscle specification (**Figure 6E)**. This pattern of *Ret*’s control of gene expression in non-neuronal tissue in the gut is even more pronounced at E14.5 where along with the musculature, genes controlling the vasculature are affected **(Figure 6H)**, while neurogenesis is largely spared. Thus, this pairwise analysis reveals that there is a gradual shift in the functional classes of genes affected by *Ret* loss-of-function, from its known primary role in neurogenesis to its non-autonomous effect on genes in the surrounding tissue in the developing gut, a feature missed by WGCNA analysis. The *Ret*-suppressed genes, in contrast at E12.5 and E14.5 were those primarily involved in epithelial development and tissue morphogenesis (**Figure 6F** **and** **6I**) hinting at *Ret*’s balancing role in controlling non-neuronal tissue development in the developing gut.

**Figure 6:**
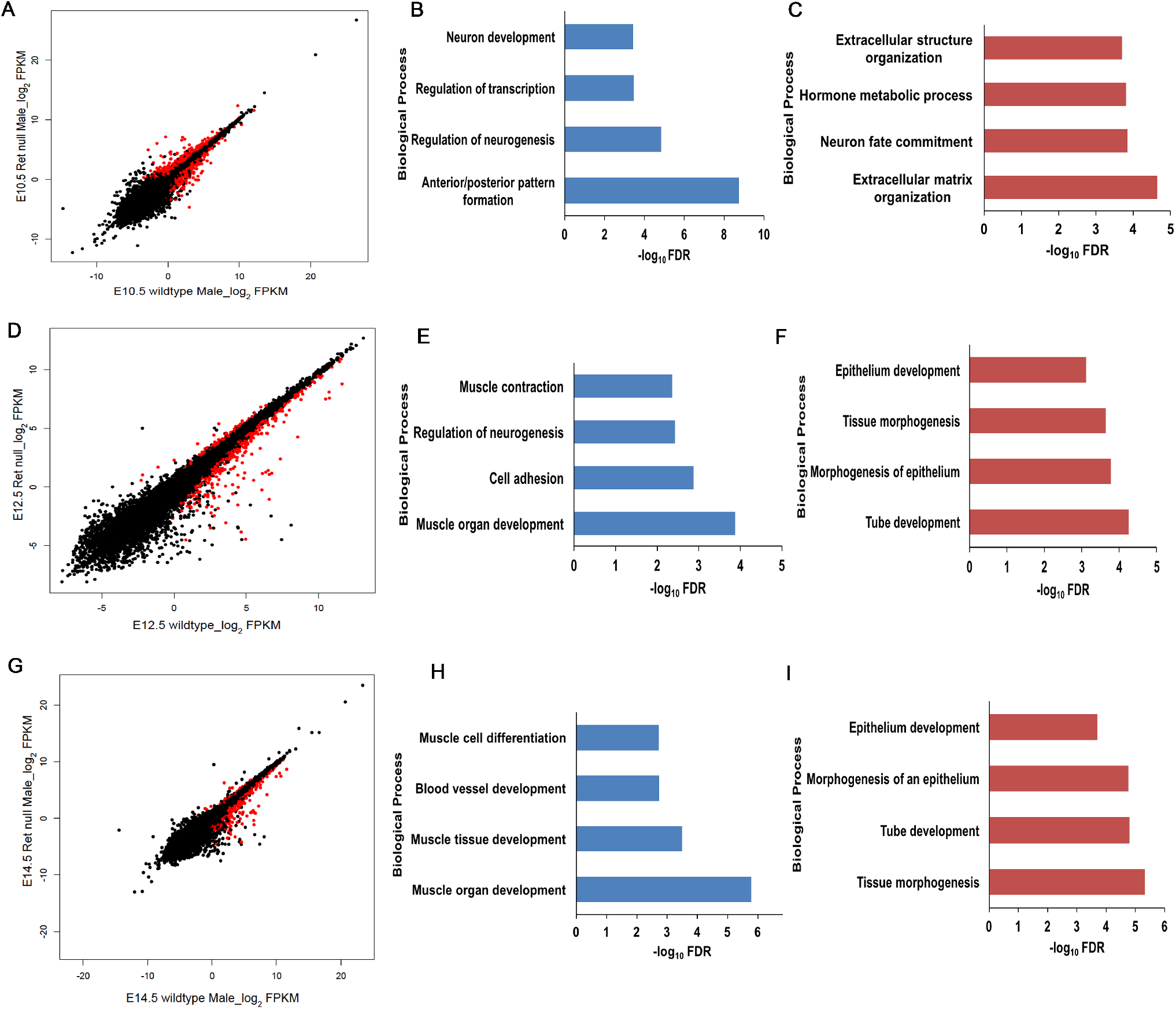
Ret-dependent gene expression in the developing gut. Scatter plots of log_2_ FPKM values between wildtype and *Ret* homozygous null embryonic gut at (A) E10.5, (D) E12.5 and (G) E14.5. Genes marked in red in the scatter plots have statistically significant (q-value <0.01) expression differences between the indicated states. Genes activated by *Ret* at E10.5 (B) includes neuronal genes and genes involved in transcriptional control, while those suppressed by *Ret* at E10.5 (C) include genes involved in extra cellular matrix formation and in hormonal response. Similar annotation analyses at E12.5 shows that along with neuronal genes, *Ret* also activates genes expressed in the musculature (E) and suppresses genes involved in epithelial development (F). At E14.5, *Ret* activates muscle-specific and vasculature development genes (H) while suppressing genes involved in epithelial morphogenesis (I).

These results show that *Ret* has a surprisingly broad cell autonomous role within enteric neurons (26) but even broader non-autonomous roles on the mesenchyme, through which the ENCCs migrate, and the vasculature, which nourishes the tissues and the epithelium in the developing gut. These non-neuronal functions are first observed at E12.5 and persist through E14.5. Further, *Ret* appears to primarily be a developmental activator since loss of *Ret* expression largely leads to transcriptional down-regulation. Thus, the two major transcriptional programs to emerge from these analyses are: (1) the predominance of multiple classes of transcription factors early in gut development gut and their control by *Ret*; and, (2) *Ret*’s cell non-autonomous role in controlling the development of the gut musculature, vasculature and epithelium. To get a better understanding of these discoveries, we studied them in greater detail.

### Transcription factors (TF) in Ret-driven gangliogenesis

One of the cardinal features of gut development is the preponderance of activities of TFs, chromatin modifiers and genes at E10.5:129 (85 down- and 44 up-regulated) TFs show significant expression differences with *Ret* loss-of-function (**Additional Data Table 9**) and are 20% of all TFs active at E10.5 (650 genes) involving many TF families. Of these, homeobox (41 members) and zinc finger (29 members) families dominate, of which *Gata2* and *Sox10* are already known direct activators of *Ret* (4), suggesting that others in this class may have similar function. With passing developmental time, a dramatically smaller and less-biased TF repertoire is affected: 34 TFs (18 down- and 16 up-regulated) at E12.5, and, 28 TFs (12 down- and 16 up-regulated) at E14.5 (**Additional Data Table 9**), primarily belonging to the zinc finger and T-box containing families. Interestingly, TF use also varies across gut development with some with various patterns of regulation. Thus, *Sox10, Snai1, Tbx2, Tbx3* and *Ascl1* are down-regulated while *Foxp1, Foxp2* and *Barx1* are up-regulated in in *Ret* null embryos through development. The effect of *Ret* loss-of-function leads to down-regulation of *Nfat5, Bnc2, Baz2b, Zfhx4 and Plag1* at E10.5 but their up-regulated at E12.5 and E14.5 in the *Ret* null. Finally, *Peg3* is downregulated in *Ret* null embryonic guts at E10.5 and E12.5 but upregulated at E14.5 (**Figure S2**). The direct functional consequences of each TF is thus non-constant through gut development rather than their presumed canonical use as activators or repressors of transcription.

Among the TFs identified, the *Hox* family members deserve careful scrutiny. *Hox* factors play critical roles in vertebrate body pattering along the anterior-posterior axis and disruption of these genes are known to have effects on neural tissues, the neural crest, endodermal and mesodermal derivatives (27–30). Multiple *Hox* genes are involved in patterning and morphogenesis of the mouse gut (31), especially genes of the *Hoxd* cluster, since their deletion (from *Hoxd4* to *Hoxd13*) leads to loss of the ileo-caecal sphincter and abnormalities in the pyloric and anal sphincter regions (32). Further, deletion of *Hoxd1* to *Hoxd10* (33) in the mouse leads to agenesis of the caecum. Almost 50% (20 out of 41) of the homeobox genes affected by loss of *Ret* at E10.5 are *Hox* genes (**Additional Data Table 9**), including the down-regulation of *Hoxd* cluster genes (*Hoxd4, Hoxd8, Hoxd9, Hoxd10, Hoxd11*, and *Hoxd13)*. Thus, *Ret* loss-of-function, with consequent induced loss of *Hoxd* gene expression, may be the causal route to aganglionosis and other gut phenotypes. The literature shows that higher *Hoxa4* expression, driven by ectopic expression in a much broader domain, leads to megacolon in the mouse from abnormal mesodermal development (34). Interestingly, *Hoxa4* has significantly higher expression in *Ret* null embryos at E10.5, and is the likely candidate for megacolon, a major unexplained phenotype of HSCR. These results point to an expanded role of *Ret* in the morphogenesis of the gut than previously thought, its importance in non-ENS phenotypes in HSCR, and, in gastrointestinal disorders generally.

### Non-autonomous effects of the ENS and the mesenchyme

When enteric neurons in the developing gut migrate through the mesenchyme, their development and activity is dependent on many trophic factors secreted by the mesenchyme, such as *Gdnf*, *Nrtn* and *Edn3,* well known for their direct effects on ENS development through their cognate receptors (18). Other factors from the surrounding epithelium like *Shh* and *Ih* also have effects on the ENS since their ablation severely reduces ENCC numbers (35, 36). Further, in rat models of HSCR, some regions of the distal colon are constricted with thick muscle and deep epithelial folds while other regions have thin external muscle with a stretched mucosa (37). This observation is assumed to be an effect of reduced enteric neuron density on the surrounding tissues, but could arise from the effects of mesenchymal defects on the ENS.

Clearly ENS defects affect the mesenchyme. Consider that although at E10.5 there is no significant enrichment of a muscle-expressing genetic program, 11 muscle-specific genes are significantly down-regulated in *Ret* null embryos at this early stage (*Tnnc2, Myh3, Myl1, Myog, Acta1, Tnnc1, Mylpf, Myl4, Actc1, Actn2, Mybpc1*). By E12.5, 18 muscle-expressed genes (*Myl6, Actc1, Cryab, Tbx2, Tbx5, Tnc, Eln, Myh6, Tagln2, Tagln3, Tnni1, Tgfb2, Tnnt2, Gpx1, Des, Meox2, Pln, Unc45b*) and by E14.5 21 muscle-expressed genes (*Myl6, Actc1, Acta1, Cryab, Tbx2, Tnc, Tbx5, Eln, Myh6, Tagln2, Ttn, Tagln3, Tnni1, Tgfb2, Tnnt2, Gpx1, Myo18b, Des, Meox2, Pln, Unc45b*) are significantly down-regulated in *Ret* null embryos and are detectably enriched as a class. The genes affected at E12.5 and E14.5 are near identical, implying a persistent, long-term mesenchymal effect from *Ret* loss-of-function. We suspected that this muscle-expressing genetic program has feedback on the ENS.

Consider that *Myog*, a transcription factor so far implicated only in the maturation of skeletal muscle (38), shows transitory expression at E10.5 in the wildtype gut but is significantly down-regulated with loss of *Ret* at E10.5. We postulated that this expression was specific to smooth muscle cells of the gut. For confirmation, we FACS-sorted EGFP-marked smooth muscle cells from the gut of a *Mhy11*-EGFP transgenic mouse at E10.5 with EGFP expression driven by a 16kb promoter sequence of the smooth muscle marker *Myh11* (39). Taqman qPCR analysis of RNA extracted from these sorted cells show that they express *Myh11*, as expected, but also *Myog,* albeit weakly, along with other smooth muscle markers, such as *Myl1* and *Myh3*, confirming the presence of *Myog* in early smooth muscle cells of the gut (**Figure 7A**). We next asked whether loss of *Myog* would lead to any ENS defects using the zebrafish, a model system known to produce aganglionosis under knockdown of *Ret* and other HSCR genes (40). Since *Myog* is expressed at a reduced level and not completely absent in *Ret* null guts we simulated this effect in zebrafish using two splice-blocking morpholinos against the intron 2-exon3 junction of *Myog*, and marked post-mitotic neurons in the developing gut with antibodies against neuronal HuC/HuD antigens to quantify the numbers of neurons (41). At 4 days post fertilization (dfp), with morpholino 1 (MO1), *myog* morphants showed curved bodies along the anterior-posterior axis, as expected, a hallmark of skeletal muscle defects. Additionally, they also demonstrated a severe loss of migratory enteric neurons as compared to embryos injected with control morpholinos (P=6.5×10^−9;^ **Figure 7B**). The anterior-posterior axis curvature was much less pronounced in morpholino 2 (MO2) injected embryos but loss of enteric neuronal precursors was equally severe (P=7.8×10^−9;^ **Figure 7B**). Thus, broader defects within the mesenchyme can also affect the migration of enteric neurons, emphasizing a two-way crosstalk between the ENS and mesenchyme necessary for normal ENS gangliogenesis.

**Figure 7:**
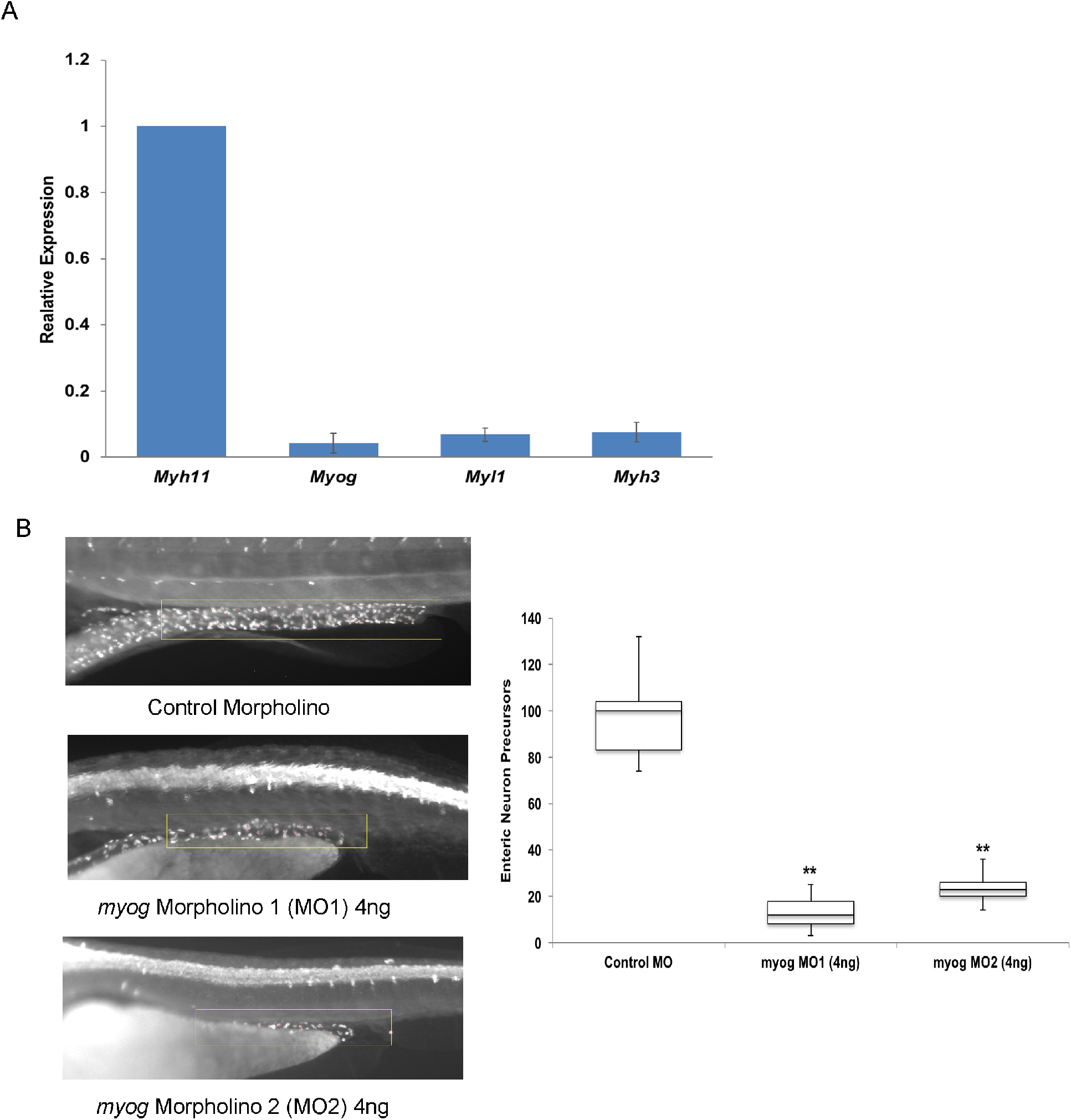
Crosstalk between enteric neurons and its supporting mesenchymal cells. (A) Gene expression qPCR of specific muscle markers were performed using FACS-sorted *Mhy11*-positive cells from *Myh11*-EGFP transgenic mouse guts at E10.5. Low level expression of *Myog*, *Myl1* and *Myh3* is observed in the gut smooth muscle cells at this stage of development. (B) Morpholino knockdown of *myog* in zebrafish leads to a skeletal muscle phenotype (significant body curvature along the anterior-posterior axis). Morpholino knockdown of *myog* in zebrafish leads to the skeletal muscle phenotype of significant body curvature along the anterior-posterior axis along with significant loss of enteric neuron precursors at 4 days post fertilization (dpf). The neuronal numbers, counted in a region of interest starting at the 8th somite and measuring till the end of the gut tube (yellow box), upon sampling 50 embryos each from wildtype and morphant embryos are significantly different (**p<0.001).

## Discussion

The gastrointestinal tract is evolutionarily one of the most primitive organs in the animal kingdom and a rudimentary form is seen in Coelenterates. In the mouse and the human, the gut tube is formed from the primitive endodermal sheet, after a series of morphogenetic movements, followed by the formation of the muscularis layers and the submucosal connective tissue from the mesoderm. Finally, neural crest-derived enteric neural crest cells (ENCC) migrate into the forming foregut to innervate the submucosal and myenteric plexus as they form throughout the gut tube, moving in a cranio-caudal manner. In the mouse, this innervation starts at E10.5 and is complete by E14.5, which is why we chose this period for developmental transcription profiling. After completion of neurogenesis at E14.5, the gut is still not fully physiologically functional and further development occurs, including after birth and consequent to feeding. However, these other time periods are less relevant to understanding human congenital motility disorders. The studies described here show that the primary feature of the E10.5 to E14.5 genetic programs is the shift from an early burst of transcription factor gene expression, indicative of the generation of cellular diversity by differentiation, to protein translation, indicating the emergence of new functions in these diverse cells, to a wider muscle, epithelium and endothelial developmental program. These trajectories are important for identifying candidate genes for various gut disorders and understanding why certain mutations are more detrimental than others (e.g., occurring earlier than later).

This study also shows that Ret signaling plays a critical and broad role in gut development in three major processes: engaging the extensive early regulatory machinery (TF), initiating ENS development and inducing neuroblast-mesenchymal interactions. A particularly surprising finding is the feedback effect of *Ret* on all three of these processes, some of which are direct and cell-autonomous (e.g., on its own regulator *Sox10*) while others are indirect and non-autonomous (e.g., on its ligand *Gdnf* or the mesenchymal TF *Myog*). Thus, the phenotypic consequences of a *Ret* deficiency are much broader than its effects on neurogenesis. These findings have three corollaries. First, HSCR patients should routinely be examined by a clinical geneticist who are likely to recognize the many additional phenotypic manifestations in HSCR beyond aganglionosis; indeed, ~32% of HSCR patients have developmental anomalies beyond aganglionosis (12). Second, the HSCR gene universe is considerably larger than currently known or assumed, including genes affecting intestinal smooth muscle and epithelium development. Third, we should not view HSCR as a cell autonomous disorder of the ENS but a multifactorial disorder of the gut involving different cell types.

These studies also provide two important lessons for the structure and functions of “systems” guiding organ development, and, therefore, human disease. As first propounded by Eric Davidson and colleagues (1, 2), development is organized around modular gene regulatory networks (GRN) that provide robustness to the system by buffering developmental noise, epigenetic noise and the effects of deleterious mutations. Thus, we need to convert our gene lists not only by functional annotations but into GRNs that *integrate* the genetic and developmental information. These GRNs are the basic ‘units’ of disease, as we have shown for the *RET* GRN and HSCR (4). Second, GRNs can display instability as well when deleterious mutations occur at certain rate-limiting genes within the GRN and lead to disease by propagating the local biochemical disruption throughout the GRN, thereby amplifying a mutational effect. We hypothesize that this feature partially explains why *RET* is the major gene for HSCR.

One remaining mystery is that despite the major sex effect in HSCR, gut development is overall not sex-biased. However, a small group of 43 genes show sex-differences in gene expression, the majority up-regulated in females and involved in diverse physiological processes. Thus, disruptions in the functions of these genes may have a larger genetic effect in males than females by virtue of the females’ higher expression, particularly if expression thresholds are important in development. These genes should be investigated for mutations in HSCR. Nevertheless, we likely have an incomplete picture of the nature and magnitude of sex differences in gut development. The mouse model we used harbors a null allele with no sex difference when aganglionosis is the measured phenotype (10). In contrast, other mouse models segregating a *Ret* null allele with a hypomorphic *Ednrb* allele do show strong sex-differences in aganglionosis and segment length (19). Thus, studying other phenotypic manifestations in mouse models may be important, for example, by quantifying the numbers and distributions of enteric neurons in the ENS. Finally, the effect of sex on transcription might not be on the whole gut or the entire ENS but on a specific group of cells, an aspect which would not be captured in our bulk transcriptomic assays but require single cell transcription studies.

## Materials and Methods

### Mouse strains

Mice heterozygous for the conditional *Ret* allele (*Ret*^fl/+;^ (42, 43), were obtained from David Ginty (Harvard University, Boston, MA). Briefly, the floxed *Ret* locus (*Ret*^fl)^ was generated by inserting a gene cassette, comprising the floxed human *RET*9 cDNA with the SV40 intron polyA sequence followed by a cyan fluorescent protein (CFP) cDNA with polyA sequence, into exon 1 of the mouse *Ret* gene; this was functionally equal to the wild-type allele (42). *Ret CFP* knock-in mice (*Ret*^CFP/+)^ were generated by crossing *Ret*^fl/+^ mice to β-actin-Cre mice to remove the *RET9* cDNA. This mouse was backcrossed >20 generation to C57BL/6 to maintain the line and all experiments conducted in this isogenic strain. The smMHC/Cre/ EGFP mice derived on a C57BL/6 background were obtained from the Jackson Laboratory (Bar Harbor, ME)(39) and were mantained in this background by repeated backcossing for >20 generations at the time of the experiments. Briefly a 16kb promoter sequence of the *Myh11* gene was used to drive *Cre* recombinase and EGFP expression. An internal ribosome entry site (IRES) of the encephalomyocarditis virus was added upstream of the EGFP cDNA to permit translation of both *Cre* and EGFP open reading frames from one mRNA. Southern blot analysis showed a single integration site of the transgene and expression in smooth muscle tissues of the developing gut (44). All animal research was approved by the Institutional Animal Care and Use Committee of the Johns Hopkins University School of Medicine.

### Mouse breeding, dissection and genotyping

Homozygous *Ret* null mice are neonatal lethals, so *Ret* heterozygous mice were intercrossed to generate all possible genotypes. Routine mouse genotyping was performed by PCR with the following primer pairs and amplicons: for the wildtype *Ret* locus, RetF 5’-CAGCGCAGGTCTCTCATCAGTACCGCA-3’ and RetR 5’-CAGCTAGCCGCAGCGACCCGGTTC-3’ resulting in a 449bp PCR product (slightly modified from (45)); for detecting the CFP knock-in we used the primer pairs *Ret*^CFP^ F 5’ ATGGTGAGCAAGGGCGAGGAGCTGTT-3’ and *Ret*^CFP^ R 5’-CTGGGTGCTCAGGTAGTGGTTGTC-3’ to amplify a 615bp PCR product from *Ret*^CFP/+^ and *Ret*^CFP/CFP^ embryos. For the Myh11-Cre-EGFP mice we used the primer pairs CreF 5’ ATGTCCAATTTACTGACCG-3’ and CreR 5’ CGCGCCTGAAGATATAGAAG-3’ to amplify a 241bp amplicon. The sex of the embryos was determined by PCR with the following primers: *Kdm5* forward 5’-CTGAAGCTTTTGGCTTTGAG-3’ and *Kdm5* reverse 5’-CCGCTGCCAAATTCTTTGG-3,’ mapping to exons 9 and 10 of the *Kdm5c/d* genes and resulting in two 331bp X chromosome-specific bands in females and an additional 301bp Y chromosome-specific band in males (46). Mouse embryos were genotyped from yolk-sac genomic DNA to allow genotype-specific analyses.

We dissected the complete gut tube at E10.5 and the developing gut from the stomach to intestine at E12.5 and E14.5 in ice cold Leibovitz’s L-15 Medium (Thermo Fischer Scientific) from wild type and *Ret*^CFP/CFP^ embryos under a fluorescence stereo zoom microscope (Zeiss AxioZoom.V16) with a suitable filter to detect CFP. The tissues were immediately flash frozen in liquid nitrogen for RNA extraction. The smMHC/Cre/ EGFP guts were dissected under a GFP filter before dissociation and cell sorting.

### Cell Sorting

Gut tubes from Myh11-Cre-EFFP embryos at E10.5 were dissociated into single cell using Accumax (Sigma, USA) by repeated pipetting. The cells were filtered serially through a 100μM and 40μM cell strainer, and centrifuged at 2000 rpm for 5 min. The cell pellet was re-suspended in 5% FBS, 4 mM EDTA in Leibovitz L-15 medium for cell sorting using MoFlo XDP (BD Biosciences) using a 488nm laser for EFGP.

### mRNA sequencing using RNA-seq

Total RNA was extracted from each of 3 male and 3 female mouse guts at E10.5, E12.5 and E14.5 from both wildtype and *Ret*^CFP/CFP^ embryos, using TRIzol (Life Technologies, USA) and cleaned on RNeasy columns (Qiagen, USA). At E10.5, owing to the small size of the embryos, two each were pooled to create one biological replicate and three replicates used. Sample integrity (>9 RIN) was assessed using an Agilent 2100 Bioanalyzer (AgilentTechnologies) and cDNA were prepared using oligo dT beads to select mRNA from total RNA followed by heat fragmentation and cDNA synthesis, as part of the Illumina Tru Seq™ RNA Sample Preparation protocol. The resultant cDNA was then used for library preparation (end repair, base ‘A’ addition, adapter ligation, and enrichment) using standard Illumina protocols. Libraries were run on a HiSeq 2000 instrument to a depth of 15 million reads per samples (75 base pair, paired end), using the manufacturer’s protocols. The primary data were generated and analyzed using the Broad Institute’s Picard Pipeline, which includes de-multiplexing and data aggregation. The resultant reads were mapped to the mouse reference genome (mm10/GRCm38) using *Tophat2* (47) with its setting for paired end, non-strand specific libraries. Successfully mapped reads were used to assemble transcripts and estimate their abundance using *Cufflinks 2.2.1* (48). The resultant data assigned Fragments Per Kilobase of transcript per Million mapped reads (FPKM) values for each transcript and gene. The transcript file for each replicate were merged using *Cuffmerge* and analyzed by *Cuffdiff* (48) to detect differentially expressed genes between various comparisons. For the purpose of this study we considered all genes with FPKM > 5 as expressed. All RNA-seq data have been deposited into NCBI’s Gene Expression Omnibus and are accessible at GEO Series accession number GSE103070.

### Weighted Gene Co-Expression Network Analysis (WGCNA)

We identified modules of co-expressed genes using the Weighted Gene Co-Expression Network Analysis (WGCNA) R package (20). A total of 36 samples were analyzed comprising 3-time points (E10.5, E12.5, and E14.5), 2 sexes (female, male), 2 *Ret* genotypes (*Ret*^+/+^, *Ret*^CFP/CFP)^ and 3 replicates each. Gene FPKM values were used for analyses after removing those with low quality in any sample (LOWDATA, HIDATA, or FAIL). There were 9,327 genes with a mean FPKM ≥ 5 across all samples retained for clustering, with expression values transformed to log_2_(FPKM+1).

We used WGCNA to conduct signed network analysis to construct modules of co-expressed genes, selecting a soft-thresholding power β of 24, the first power with adjusted R^2^ > 0.8, to satisfy the assumption of a scale-free topology (20). We used the “bi-weight mid-correlation” correction method, instead of the Pearson correlation coefficient, because it has improved performance when outliers are present. We set the parameter pamRespectsDendro to TRUE, and, to obtain smaller, more numerous modules, we set the minimum module size to 25 and the deepSplit parameter value to 2. We tested association of each module eigengene, which is the first principal component and represents the module’s expression profile, with each of five binary variables using linear regression: genotype (*Ret*^+/+^, *Ret*^CFP/CFP)^; sex (males vs. females); E10.5 vs. non-E10.5; E12.5 vs. non-E12.5; E14.5 vs. non-E14.5. Module associations with p-values below the Bonferroni-adjusted threshold, accounting for the number of modules, were considered statistically significant.

### Taqman gene expression assays

Total RNA was extracted from pooled EGFP FACS-sorted cells from the guts of 3 smMHC/Cre/EGFP embryos at E10.5 using TRIzol (Life Technologies, USA) and cleaned on RNeasy columns (Qiagen, USA). Sample integrity was assessed using an Agilent 2100 Bioanalyzer (AgilentTechnologies) and 100ng of total RNA was converted to cDNA using SuperScriptIII reverse transcriptase (Life Technologies, USA) and Oligo-dT primers. The total cDNA was subjected to Taqman gene expression analyses (Life Technologies, USA) using transcript-specific probes and primers. For the whole gut experiments to titrate the correlation between FPKM and Ct, 500 ng of RNA from each biological replicate was used to make cDNA. Mouse β-actin (*Actb*) was used as an internal loading control for normalization. Each assay was performed in triplicate (n=9 observations per state); the data are presented as means with their standard errors. Relative fold changes were calculated based on the 2^ΔΔCt^(threshold cycle) method for measuring gene expression, the value for the gene with the highest expression (lowest Ct value) being set to unity. The following mouse Taqman probes were obtained from Applied Biosystems: *Syt6* (Mm00490071_m1), *Trpc6* (Mm01151079_m1), *Mnx1* (Mm00658300_g1), *Tead4* (Mm01189836_m1), *Dnajc12*(Mm00497038_m1), *Plekhb2* (Mm01234649_m1), *Cdh23* (Mm04335689_g1), *Kif5a* (Mm00515265_m1), *Angpt4* (Mm00507766_m1), *Myh11 (*Mm00443013_m1*), Myog (*Mm00446194_m1*), Myl1 (*Mm00659043_m1) and *Myh3(*Mm01332463_m1).

### Zebrafish maintenance, embryo collection and morpholino injections

Zebrafish (AB strain) were raised and maintained under standard conditions, with embryos collected and staged as previously described (49). All animal research was approved by the Institutional Animal Care and Use Committee of the Johns Hopkins University School of Medicine.

For zebrafish gene knockdown experiments, two splice blocking morpholinos (MO) were designed at the intron 2-exon 3 junction of *myog;* MO1: 5’-ATCTGAGAAAAGTAGACCACAATGT-3’ and MO2: 5’-GACGACACCTGTCACACACACAC-3’ (Gene Tools, LLC), along with a standard negative control morpholino (5’-CCTCTTACCTCAGTTACAATTTATA-3’) targeting a human beta-globin intron mutation for beta-thalassemia. We also injected a p53 morpholino (5’-GCGCCATTGCTTTGCAAGAATTG-3’) at 2ng concentration along with our transcript-specific morpholinos because this morpholino has been reported to suppress the apoptotic effects induced by some morpholinos (50). Injections were performed on ~30 one-to two-cell zebrafish embryos independently on at least 3 different days (n=100). Different concentrations (from 1 to 6ng) of MO were injected to determine the optimal concentration at which a phenotype could be consistently detected. The data shown are for 4ng where there was a measurable phenotype for enteric neurons at 4 dpf (days post-fertilization) without severe malformation or death of embryos.

### Immunostaining, visualization and cell counting

Injected zebrafish embryos were fixed at 4 dpf with 4% paraformaldehyde (PFA) for 2 hours at room temperature. A monoclonal anti-HuC antibody (Invitrogen #A-21271), followed by Alexa Fluor 568 F (ab’)_2_ fragment of goat anti-mouse IgG secondary antibody (Invitrogen #A11019), was used for fluorescent labeling of enteric neurons as previously described (40, 51), with mild modifications. The embryos were bleached after fixing in 4% PFA by incubating them in 3% H_2_O_2_/0.5% KOH medium until there was complete loss of epidermal pigmentation (~20 min), followed by a 5 min wash with PBS to stop the bleaching reaction. 50 stained embryos (for both wildtype and morphants) were visualized using a Zeiss AxioZoomV16 fluorescent microscope using a DS red filter to assess the colonization of enteric neurons in the gut of each embryo. Stained neurons were counted using the Image-based Tool for Counting Nuclei (ITCN) plugin in ImageJ (41), with the following parameters: width 9 pixels, minimum distance 4.5 pixels, threshold of 1 and using a selected Region of Interest (ROI). Since enteric neurons in Hirschsprung disease models in zebrafish are mostly lost caudally in the gut, we chose our ROI as 8 somites starting at the caudal end of the gut and moving rostrally. At least 20 embryos were used for cell counting for each concentration of morpholino used. P values were calculated from a pairwise 2-tailed t-test.

## AUTHOR CONTRIBUTIONS

SC and AC conceived and designed the study. SC and DRA conducted all embryonic dissections, tissue collection and molecular biology. SC performed all *in vitro* and zebrafish transgenic assays. SBG conducted RNA-seq experiments. PN and SC performed the RNA–seq analysis. SC and AC wrote the manuscript. All authors contributed to manuscript editing. These studies were supported by an NIH MERIT Award HD28088 to A.C.

## References

1. Davidson EH (2006) The regulatory genome: gene regulatory networks in development and evolution (Academic, Burlington, MA) New Ed pp xi, 289 p.

2. Davidson EH (2010) Emerging properties of animal gene regulatory networks. Nature 468(7326):911–920.

3. Chakravarti A & Turner TN (2016) Revealing rate-limiting steps in complex disease biology: The crucial importance of studying rare, extreme-phenotype families. Bioessays 38(6):578–586.

4. Chatterjee S, et al. (2016) Enhancer Variants Synergistically Drive Dysfunction of a Gene Regulatory Network In Hirschsprung Disease. Cell 167(2):355–368 e310.

5. Furness JB (2006) Structure of the Enteric Nervous System. (Blackwell Publishing, Malden, Massachusetts, USA), pp 1–28.

6. Heanue TA & Pachnis V (2007) Enteric nervous system development and Hirschsprung’s disease: advances in genetic and stem cell studies. Nat Rev Neurosci 8(6):466–479.

7. Shivdasani RA (2002) Molecular regulation of vertebrate early endoderm development. Dev Biol 249(2):191–203.

8. de Santa Barbara P, van den Brink GR, & Roberts DJ (2003) Development and differentiation of the intestinal epithelium. Cell Mol Life Sci 60(7):1322–1332.

9. Chakravarti A MAS, Lyonnet S (2001) Hirschsprung Disease. The Metabolic and Molecular Bases of Inherited Disease, ed Valle D BAL, Vogelstein B, Kinzler K.W., Antonarakis S.E., Ballabio A, Gibson K, Mitchell G (McGraw-Hill, New York), 8 Ed.

10. Uesaka T, Nagashimada M, Yonemura S, & Enomoto H (2008) Diminished Ret expression compromises neuronal survival in the colon and causes intestinal aganglionosis in mice. J Clin Invest 118(5):1890–1898.

11. Arighi E, Borrello MG, & Sariola H (2005) RET tyrosine kinase signaling in development and cancer. Cytokine Growth Factor Rev 16(4-5):441–467.

12. Tilghman JM, et al. (2019) Molecular Genetic Anatomy and Risk Profile of Hirschsprung’s Disease. N Engl J Med 380(15):1421–1432.

13. Schuchardt A, D’Agati V, Larsson-Blomberg L, Costantini F, & Pachnis V (1994) Defects in the kidney and enteric nervous system of mice lacking the tyrosine kinase receptor Ret. Nature 367(6461):380–383.

14. Gerrard DT, et al. (2016) An integrative transcriptomic atlas of organogenesis in human embryos. Elife 5.

15. Kukurba KR & Montgomery SB (2015) RNA Sequencing and Analysis. Cold Spring Harb Protoc 2015(11):951–969.

16. Pervouchine DD, et al. (2015) Enhanced transcriptome maps from multiple mouse tissues reveal evolutionary constraint in gene expression. Nat Commun 6:5903.

17. Heanue TA & Pachnis V (2006) Expression profiling the developing mammalian enteric nervous system identifies marker and candidate Hirschsprung disease genes. Proc Natl Acad Sci U S A 103(18):6919–6924.

18. Obermayr F, Hotta R, Enomoto H, & Young HM (2013) Development and developmental disorders of the enteric nervous system. Nat Rev Gastroenterol Hepatol 10(1):43–57.

19. McCallion AS, Stames E, Conlon RA, & Chakravarti A (2003) Phenotype variation in two-locus mouse models of Hirschsprung disease: tissue-specific interaction between Ret and Ednrb. Proc Natl Acad Sci U S A 100(4):1826–1831.

20. Langfelder P & Horvath S (2008) WGCNA: an R package for weighted correlation network analysis. BMC Bioinformatics 9:559.

21. Huang da W, Sherman BT, & Lempicki RA (2009) Systematic and integrative analysis of large gene lists using DAVID bioinformatics resources. Nat Protoc 4(1):44–57.

22. Le Douarin NM & Teillet MA (1973) The migration of neural crest cells to the wall of the digestive tract in avian embryo. J Embryol Exp Morphol 30(1):31–48.

23. Emison ES, et al. (2010) Differential contributions of rare and common, coding and noncoding Ret mutations to multifactorial Hirschsprung disease liability. Am J Hum Genet 87(1):60–74.

24. Emison ES, et al. (2005) A common sex-dependent mutation in a RET enhancer underlies Hirschsprung disease risk. Nature 434(7035):857–863.

25. Kapoor A, et al. (2015) Population variation in total genetic risk of Hirschsprung disease from common RET, SEMA3 and NRG1 susceptibility polymorphisms. Hum Mol Genet 24(10):2997–3003.

26. Durbec PL, Larsson-Blomberg LB, Schuchardt A, Costantini F, & Pachnis V (1996) Common origin and developmental dependence on c-ret of subsets of enteric and sympathetic neuroblasts. Development 122(1):349–358.

27. Krumlauf R (1994) Hox genes in vertebrate development. Cell 78(2):191–201.

28. Mallo M, Wellik DM, & Deschamps J (2010) Hox genes and regional patterning of the vertebrate body plan. Dev Biol 344(1):7–15.

29. Wellik DM (2009) Hox genes and vertebrate axial pattern. Curr Top Dev Biol 88:257–278.

30. Manley NR & Capecchi MR (1998) Hox group 3 paralogs regulate the development and migration of the thymus, thyroid, and parathyroid glands. Dev Biol 195(1):1–15.

31. Beck F (2002) Homeobox genes in gut development. Gut 51(3):450–454.

32. Zakany J & Duboule D (1999) Hox genes and the making of sphincters. Nature 401(6755):761–762.

33. Zacchetti G, Duboule D, & Zakany J (2007) Hox gene function in vertebrate gut morphogenesis: the case of the caecum. Development 134(22):3967–3973.

34. Wolgemuth DJ, Behringer RR, Mostoller MP, Brinster RL, & Palmiter RD (1989) Transgenic mice overexpressing the mouse homoeobox-containing gene Hox-1.4 exhibit abnormal gut development. Nature 337(6206):464–467.

35. Ramalho-Santos M, Melton DA, & McMahon AP (2000) Hedgehog signals regulate multiple aspects of gastrointestinal development. Development 127(12):2763–2772.

36. Mao J, Kim BM, Rajurkar M, Shivdasani RA, & McMahon AP (2010) Hedgehog signaling controls mesenchymal growth in the developing mammalian digestive tract. Development 137(10):1721–1729.

37. Stamp LA, et al. (2015) Surgical Intervention to Rescue Hirschsprung Disease in a Rat Model. J Neurogastroenterol Motil 21(4):552–559.

38. Sassoon D, et al. (1989) Expression of two myogenic regulatory factors myogenin and MyoD1 during mouse embryogenesis. Nature 341(6240):303–307.

39. Xin HB, Deng KY, Rishniw M, Ji G, & Kotlikoff MI (2002) Smooth muscle expression of Cre recombinase and eGFP in transgenic mice. Physiol Genomics 10(3):211–215.

40. Jiang Q, et al. (2015) Functional loss of semaphorin 3C and/or semaphorin 3D and their epistatic interaction with ret are critical to Hirschsprung disease liability. Am J Hum Genet 96(4):581–596.

41. Schneider CA, Rasband WS, & Eliceiri KW (2012) NIH Image to ImageJ: 25 years of image analysis. Nat Methods 9(7):671–675.

42. Uesaka T, Nagashimada M, Yonemura S, & Enomoto H (2008) Diminished Ret expression compromises neuronal survival in the colon and causes intestinal aganglionosis in mice. J Clin Invest 118(5):1890–1898.

43. Gould TW, Yonemura S, Oppenheim RW, Ohmori S, & Enomoto H (2008) The neurotrophic effects of glial cell line-derived neurotrophic factor on spinal motoneurons are restricted to fusimotor subtypes. J Neurosci 28(9):2131–2146.

44. Madsen CS, et al. (1998) Smooth muscle-specific expression of the smooth muscle myosin heavy chain gene in transgenic mice requires 5’-flanking and first intronic DNA sequence. Circ Res 82(8):908–917.

45. Jain S, et al. (2004) Mice expressing a dominant-negative Ret mutation phenocopy human Hirschsprung disease and delineate a direct role of Ret in spermatogenesis. Development 131(21):5503–5513.

46. Clapcote SJ & Roder JC (2005) Simplex PCR assay for sex determination in mice. Biotechniques 38(5):702, 704, 706.

47. Kim D, et al. (2013) TopHat2: accurate alignment of transcriptomes in the presence of insertions, deletions and gene fusions. Genome Biol 14(4):R36.

48. Trapnell C, et al. (2012) Differential gene and transcript expression analysis of RNA-seq experiments with TopHat and Cufflinks. Nat Protoc 7(3):562–578.

49. Kimmel CB, Ballard WW, Kimmel SR, Ullmann B, & Schilling TF (1995) Stages of embryonic development of the zebrafish. Dev Dyn 203(3):253–310.

50. Robu ME, et al. (2007) p53 activation by knockdown technologies. PLoS Genet 3(5):e78.

51. Kuhlman J & Eisen JS (2007) Genetic screen for mutations affecting development and function of the enteric nervous system. Dev Dyn 236(1):118–127.

