## Supplemental Information for "The gastrointestinal development ‘parts list’: transcript profiling of embryonic gut development in wildtype and *Ret*-deficient mice"

**This PDF file includes:**

Figs. S1to S2
Captions for datasets S1to S9

**Other supplementary materials for this manuscript include the following:**

Datasets S1to S9

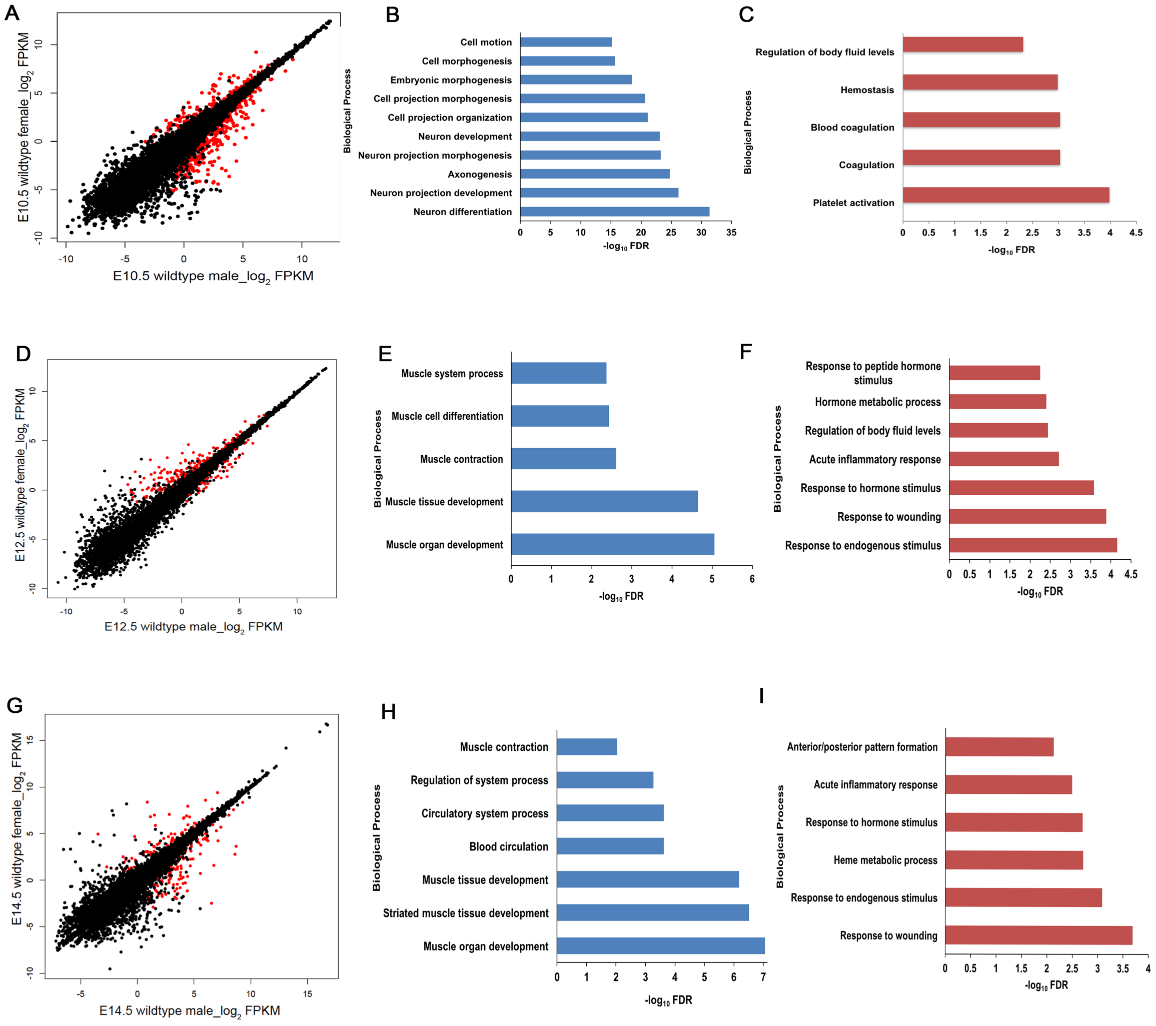


**Fig. S1**. *Sex-dependent gene expression in the embryonic mouse gut.*
(A) Scatter plot of $log_2$ FPKM values of genes expressed in wildtype males versus females at E10.5. (B) GO annotation clustering of genes with sex differences shows enrichment of cell motility and neuronal differentiation genes in males, and (C) those controlling homeostasis and blood coagulation processes in females. (D) Analogous analysis of data from E12.5 shows a higher expression of muscle specific genes in males (E) and those controlling hormonal processes and inflammatory responses in females (F). Scatter plot for analogous data at E14.5 also highlights sex-differences in gene expression (G), with higher expression of muscle specification and vasculature genes in males (H) and epithelial morphogenesis in females (I). Genes marked in red in the scatter plots have statistically significant (q-value <0.01) expression differences between the indicated states.

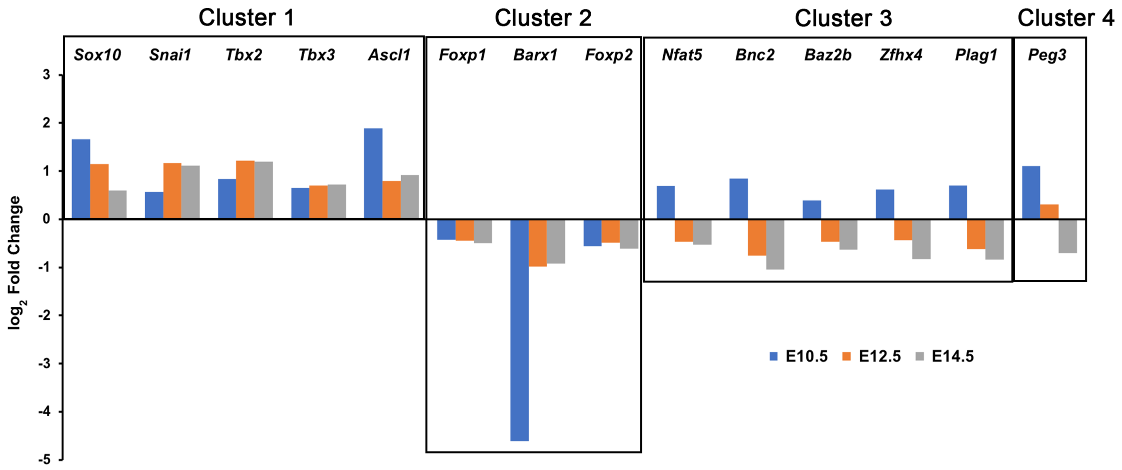


**Fig. S2**. *Differential effect of Ret loss of function of transcription factors (TFs) through development.*
The 14 transcription factors which are affected in the *Ret* null embryonic guts at all stages in development shows distinct temporal response patterns. Cluster 1 contains TFs which are downregulated all through development whereas cluster 2 contain TFs which are upregulated all through development in Ret null embryos. Cluster 3 are a group of TFs downregulated at E10.5 but upregulated at E12.5 and E14.5 null embryos. Cluster 4 which contain only one TF (Peg3) is downregulated in E10.5 and E12.5 but upregulated at E14.5. The measurement is the log2 scale of the fold change (Wildtype FPKM/ Ret null FPKM) at all 3 stages.

**Additional data table S1 (separate file)**

7793 genes with a mean FPKM ≥ 5 across all 36 samples grouped into 18 modules which were used to test associations between the first principal component (the eigengene) of each module's expression profile with each of five binary variables using linear regression.

**Additional data table S2 (separate file)**

List of genes showing significant differential expression between males and females at E10.5 mouse gut.

**Additional data table S3 (separate file)**

List of genes showing significant differential expression between males and females at E12.5 mouse gut.

**Additional data table S4 (separate file)**
List of genes showing significant differential expression between males and females at E14.5 mouse gut.

**Additional data table S5 (separate file)**
List of genes showing significant differential expression between wildtype male guts at E10.5 (early) compared to E14.5 (late).

**Additional data table S6 (separate file)**
List of genes showing significant differential expression between wildtype and Ret homozygous null mouse guts at E10.5

**Additional data table S7 (separate file)**
List of genes showing significant differential expression between wildtype and Ret homozygous null mouse guts at E12.5

**Additional data table S8 (separate file)**
List of genes showing significant differential expression between wildtype and Ret homozygous null mouse guts at E14.5

**Additional data table S9 (separate file)**
List of transcription factors showing significant differential expression between wildtype and Ret homozygous null mouse guts at E10.5, E12.5 and E14.5
